# Gut-immune reactivity underlies sex differences in the maternal immune activation preclinical model of autism

**DOI:** 10.1101/2024.06.18.599415

**Authors:** Stephanie Salia, Francine F. Burke, Meagan E. Hinks, Alison M. Randell, Mairead Anna Matheson, Susan G. Walling, Ashlyn Swift-Gallant

## Abstract

The gut microbiome plays a vital role in health and disease, including neurodevelopmental disorders like autism spectrum disorder (ASD). ASD affects 4:1 males-to-females, and sex differences are apparent in gut microbiota composition among ASD individuals and in animal models of this condition, such as the maternal immune activation (MIA) mouse model. However, few studies have included sex as a biological variable when assessing the role of gut microbiota in mediating ASD symptoms. Using the MIA model of ASD, we assessed whether gut microbiota contributes to the sex differences in the presentation of ASD-like behaviors. Gut microbiota transplantation from MIA or vehicle/control male and female mice into healthy, otherwise unmanipulated, 4-week-old C57Bl/6 mice was performed for 6 treatments over 12 days. Colonization with male, but not female, MIA microbiota was sufficient to reduce sociability, increase repetitive burying behavior, decrease microbiota diversity and increase neuroinflammation with more pronounced deficits in male recipients. Colonization with both male and female donor microbiota altered juvenile ultrasonic vocalizations and anxiety-like behavior in recipients of both sexes, and there was an accompanied change in the gut microbiota and serum cytokine IL-4 and IL-7 levels of all recipients of MIA gut microbiota. In addition to the increases in gut microbes associated with pathological states, the female donor microbiota profile also had increases in gut microbes with known neural protective effects (e.g., *Lactobacillus* and *Rikenella*). These results suggest that gut reactivity to environmental insults, such as in the MIA model, plays a pivotal role in shaping the sex disparity observed in ASD development.

**Significance Statement:** Increasing evidence suggests a role for the gut microbiota in autism spectrum disorder (ASD). ASD development has largely been associated with genetic mutations; however, with a 4-fold greater risk among males than females, the sex of a child is a predictor equivalent to familial ASD incidence. Using a preclinical mouse model of ASD, maternal immune activation (MIA), we show that gut microbiota transfer from MIA males into unaffected control males was most effective in reproducing ASD-like symptoms and led to distinct gut microbiota composition and greater inflammation in recipients. These findings suggest that how the gut responds to environmental insults differs between sexes, and this variance contributes to the greater risk ASD development among males than females.

## Introduction

Autism spectrum disorder (ASD) is a complex neurodevelopmental disorder (NDD) characterized by challenges in communication, social interactions and repetitive or restricted behaviors (1). It is estimated that 1 in 100 children is diagnosed with ASD globally (2), posing a significant economic burden for families and societies (3). ASD development has largely been associated with genetic mutations (4); however, with a 4-fold greater risk among males than females, the sex of a child is a predictor equivalent to familial ASD incidence (5, 6). As such, it is likely that sex-specific factors in early development contribute to ASD etiology (6,7).

Gestational infection, and the resulting maternal immune activation (MIA), are risk factors for ASD, and can be reliably modelled in rodents (8). MIA triggers heightened maternal pro-inflammatory cytokines and chemokines, such as interleukin (IL)-6 and IL-17α, which can interfere with fetal brain development (9,10). In rodent models, MIA is induced through gestational exposure to the viral mimic polyinosinic: polycytidylic acid (poly IC; 10). Analogous to ASD in humans, MIA male offspring are more likely than females to develop an ASD-like phenotype in this model (8,11).

Recent studies have shown that MIA leads to dysbiosis of the gut microbiota (12,13). This is significant because the gut microbiota, residing in our gastrointestinal (GI) tract, influences immune, metabolic, and nervous system development via the microbiota-gut-brain axis (14,15), potentially contributing to ASD-like traits in the MIA model. Indeed, gut microbiota transplantation from human ASD donors to mice without microbiota (i.e., germ-free) has been shown to induce ASD-like phenotypes (16,17). Although a possible association between biological sex differences and gut microbiota has been proposed, studies assessing the role of the gut microbiota in ASD development largely study only males and/or omit the sex/gender of donors and recipients. From the few studies that include sex as a biological variable, there are indications for sex-specific mediation of gut-immune effects in the MIA model. For example, behavioral changes are more pronounced in male recipient mice of human male ASD gut microbiota (16), and sex differences in gut microbiota diversity contribute to sex differences in social behavior (18–20).

There are also indications that the neuroimmune system is sexually differentiated and contributes to male vulnerability to neurodevelopmental insults. For instance, the effect of the gut microbiota on the neural immune cells, microglia, varies with sex and age, with male microglia being more sensitive to microbiota perturbation during prenatal development (21). Despite this evidence, sex is often overlooked in preclinical studies investigating how gut microbiota influences ASD/MIA symptoms, with many studies exclusively using male subjects and/or microbiota from male donors (e.g., 16,17), while others fail to report the sex of the animals used.

The present study systematically evaluated the contributions of the gut microbiota to sex differences in the presentation of ASD-like behaviors using the MIA preclinical mouse model of ASD. We transplanted same- or other-sex cecal microbiota from MIA or vehicle/control mice into healthy/otherwise unmanipulated male and female recipient mice to determine whether MIA-induced ASD-like phenotype was transmissible via the gut microbiota (Fig. 1A). We then assessed the effects of recipient sex (male or female) and donor cecum sex/treatment effects on the (i) ASD-like behavioral phenotype (ii) gut microbiota composition, (iii) circulating inflammatory cytokines, and (iv) neuroinflammatory response in the hippocampal dentate gyrus (DG). We hypothesized that sex would modulate the effects of microbiota transplantation in recipient mice, and the sex-based analyses would allow us to identify gut-immune pathways that contribute to the sex-differential vulnerability to gestation infection.

**Figure 1:**
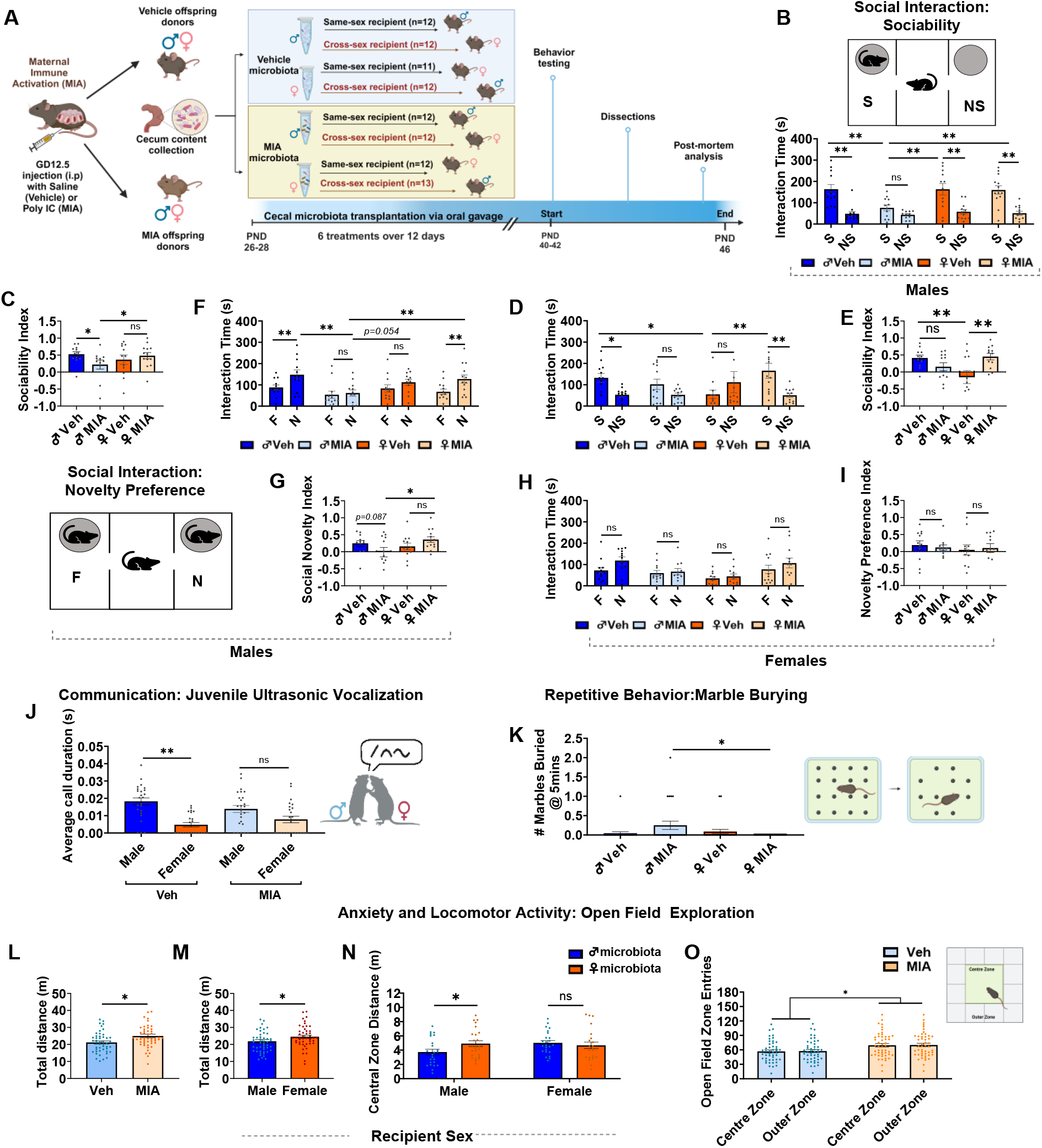
**Note:** Cross and same-sex MIA microbiota transplant promote ASD-like phenotype in WT mice. (A) Experimental design: Preparation of cecal microbiota transplantation from MIA-exposed mice prenatally treated with poly I:C (5mg/kg) or vehicle (0.9% sterile NaCl) solution on GD12.5. WT mice were colonized with 6 treatments (every other day for 12 days) with cecal microbiota from Vehicle (Veh) or MIA donors at weaning. Recipient mice were behaviorally tested starting at 6 weeks of age. (B-O) Behavioral phenotypes in recipient mice. (B-I) time interacting with social and novel stimuli in the three-chamber test. Male MIA cecum transplant led to a social and novelty preference deficit in male recipient mice but not in females. (J) MIA microbiota alters opposite-sex social communication by reducing ultrasonic vocalizations and average call duration in male recipient mice. (K) Male, but not female, donor MIA microbiota increased marbles buried at 5mins intervals in the marble burying test. (L-O) MIA cecum transplant increases locomotor/hyperactivity in male and female recipients, evidenced by distance travelled and zone entries in the open field test. Data represent the mean ± SEM (*n* = 11-13 per group; \**p* < 0.05, \*\**p* < 0.01; n.s., not significant; S, social; NS, non-social; F, familiar and N, novel stimuli. Fig.A created with Biorender.com.

## Results

### Male, but not female, MIA microbiota transplant promotes social behavioral deficit in male recipient mice

To determine the effect of MIA microbiota transplant on recipients’ social behavior, we compared WT male and female mice colonized with microbiota from either male vehicle (♂Veh), female vehicle (♀Veh), male MIA (♂ MIA), or female MIA (♀ MIA) donor mice in the three-chamber sociability test. We found that male mice that received ♂MIA microbiota spent significantly less time than male mice that received ♂Veh, ♀Veh, or ♀MIA microbiota investigating the age and sex-matched unfamiliar social stimulus mouse, and did not exhibit a significant preference for the social stimulus over the non-social stimulus (i.e., empty holding cup), *F* (1, 45(cecum sex X treatment)) = 4.22, *p* = 0.046; post hoc tests, all *p* values < 0.01, (Fig. 1B). Additionally, male recipients of ♂MIA microbiota displayed a corresponding decrease on the social preference index than recipients of ♂Veh and ♀MIA microbiota [*F* (1, 45(cecum sex X treatment)) = 4.71, *p* = 0.035; post hoc tests, all *p* values < 0.01; Fig. 1C]. Conversely, both female mice that received ♂MIA and ♀Veh microbiota did not show a significant preference for the social stimulus over the non-social stimulus, *F* (1, 43(cecum sex X treatment)) = 7.53, *p* = 0.009; post hoc tests, all *p* values < 0.05 (Fig. 1D). Time spent interacting with the social stimulus and the social preference index were also lower in female mice that received ♀Veh microbiota than female recipients of ♀MIA and ♂Veh microbiota (*p* =0.044; Fig. 1d), and *F* (1, 42(cecum sex X treatment)) = 11.34, *p* = 0.002; post hoc tests, all *p* values < 0.01; Fig. 1E, respectively. This unexpected finding suggests that male donor microbiota may increase social behavior in female recipients. Overall, ♂MIA microbiota had a minimal effect on the social behavior of female recipient mice.

Similar results were found on the social novelty preference trial, where experimental mice were simultaneously exposed to a familiar and novel social stimulus mouse. We found that male recipients of ♂MIA microbiota spent significantly less time interacting with the novel mouse and did not display a clear preference for the novel over the familiar mouse [(*F* (1, 45) (cecum sex X treatment) = 5.28, *p* = 0.026, post hoc tests, all *p* values < 0.05; Fig. 1F]. Male recipients of ♀Veh microbiota also did not show a preference for the novel stimulus suggesting marginal deficits on social novelty preference (Fig. 1F). Further, male recipients of ♂MIA microbiota showed a significantly lower social novelty preference index when compared to male recipients of ♂Veh microbiota [(*F* (1, 45) (cecum sex X treatment) = 5.01, *p* = 0.030, post hoc tests, *p* = 0.087; Fig. 1G]. Female recipient mice showed no difference in time spent exploring the novel versus familiar stimulus and correspondingly did not exhibit a difference in the novelty preference index (all *p* values > 0.100; Fig. 1H-1I). Together, the results indicate that male MIA microbiota transplant alters social behavior and social novelty preference in males than in female recipient mice when compared to their same-sex control groups.

### Social communication is altered with MIA microbiota transplant

Social communication was assessed via juvenile ultrasonic vocalizations (USVs) production during social interaction with unfamiliar male and female conspecifics (22). We observed a significant interaction between recipient sex and cecum treatment on the average call duration during social interaction with an unfamiliar female stimulus [*F* (1, 88)(sex X cecum treatment) = 4.01, *p* = 0.048]. While the predicted sex difference was found such that male recipients of vehicle microbiota (regardless of donor sex) made longer calls than female recipients (vehicle or MIA microbiota) (*p* < 0.001), no sex difference was found among recipients of MIA microbiota; male recipients of MIA microbiota did not differ from female recipients of MIA microbiota (*p* > 0.100; Fig. 1J). Overall, we found the expected recipient main effect of sex (males > females) on all USV parameters during social interaction with an unfamiliar female stimulus. Specifically, regardless of cecum sex or treatment, male recipient mice made more calls and called for longer duration and at higher frequencies than female recipient mice (all *p* values < 0.001; SI Table 1). In contrast, no differences were found in the total number of calls, total call duration, average call duration, and mean low and high frequency of calls produced during interaction with an unfamiliar male mouse (all *p* values > 0.100). These results are consistent with previous reports of sex differences in USV call duration (23) and suggest MIA microbiota eliminates this sex difference, which may have implications for socio-sexual behaviors in rodents (20).

### Repetitive behavior is increased with male-MIA microbiota transplant

The marble bury test was used to measure the tendency of mice to engage in digging behavior; while digging is a natural behavior, excessive digging and burying are representative of repetitive behavior (24). Significant cecum sex by cecum treatment interaction was found in the number of marbles buried at 5 min [*F* (1, 88) = 5.62, *p* = 0.020]. ♂MIA microbiota transplant promoted repetitive burying behavior in both male and female recipients compared to ♀MIA microbiota (i.e., data combined) as demonstrated by increases in marbles buried compared to veh controls (*p* = 0.029; Fig. 1K). These results indicate that male-specific gut microbes promote repetitive behavior in both male and female mice.

### MIA microbiota transplant increases locomotor/hyperactivity in male and female recipients

Locomotor activity and anxiety-like behavior, common comorbidity of ASD, were assessed using the open field test (25, 26). On the total distance travelled, we found a significant main effect of recipient sex, [*F* (1, 88) = 3.99, *p* = 0.049], such that, regardless of cecum sex or treatment, female recipient mice travelled more distance than male recipients in the open field arena (p = 0.049; Fig. 1L). A significant main effect of cecum treatment was also found on total distance travelled [*F* (1, 88) = 7.75, *p* = 0.007), with recipients of MIA microbiota transplant, regardless of donor cecum sex, travelling longer distance than vehicle microbiota recipient mice (*p* = 0.007; Fig. 1M), indicating increased locomotor activity. Time spent in the central and outer zones of the open field showed no significant differences for all groups (all *p* values > 0.100). However, we found a significant recipient sex-by-cecum sex interaction on central zone distance travelled [*F* (1, 88) = 3.96, *p* = 0.049] such that male recipients of male microbiota, regardless of treatment, travelled less distance in the center zone than male recipients of female microbiota or female recipients of male microbiota, suggestive of greater anxiety-like behavior in male recipients of male microbiota (all *p* values < 0.05; Fig. 1N). Additionally, MIA microbiota transplant, regardless of donor cecum sex, increased the number of entries into the center and outer zones in both male and female recipients, compared to vehicle-microbiota recipient mice (all *p* values < 0.05; Fig. 1O), suggesting that MIA microbiota increased locomotor both male and female recipients.

### MIA microbiota transplant, regardless of donor sex, alters gut microbiota composition in recipient mice

To evaluate whether microbiota transplant from male and female MIA donors induces changes in recipients gut microbial composition, we examined cecal contents by 16S rRNA gene sequencing of samples isolated from male and female mice that received either vehicle (i.e., ♂ or ♀) or MIA (i.e., ♂ or ♀) microbiota transplant. Overall, *Bacteroidetes* and *Firmicutes* were the dominant phyla in the microbiota composition of all groups (Fig. 2A); however, the *Firmicutes/Bacteroidetes* ratio, which has been reported to be higher in MIA-treated offspring (27) showed no difference across all groups (*p* > 0.100). Taxonomic analyses revealed that phyla *Deferribacterota* and *Actinobacteriota*, the families of *Ruminococcaceae, Deferribacteraceae, Erysipelotrichaceae* and *Anaerovoracaceae,* and the genera of *Blautia*, *Tuzzerella, Lachnospiraceae, Eubacterium* and *Mucispirillum* had significantly lower relative abundances in mice that received MIA microbiota, regardless of donor or recipient sex. In contrast, the relative abundance of the phyla *Proteobacteria*, families of *Butyricicoccaceae* and *Sutterellaceae,* and the genera of *Prevotellaceae, Odoribacter, Rikenella,* and *Parasutterella* were significantly higher in MIA microbiota recipient mice (all *p* values < 0.05; Table 1). The abundance of *Lachnospiraceae*, which has been correlated with greater social behavior deficit in children with ASD (28), was significantly higher in male mice that received MIA microbiota (*p* < 0.05; Fig. 2B). These differences in the gut microbial community indicate that specific microbial taxa are altered in a sex-dependent manner with MIA microbiota transplant and may contribute to the ASD-like behavioral deficits observed in the MIA microbiota recipient mice.

**Figure 2.**
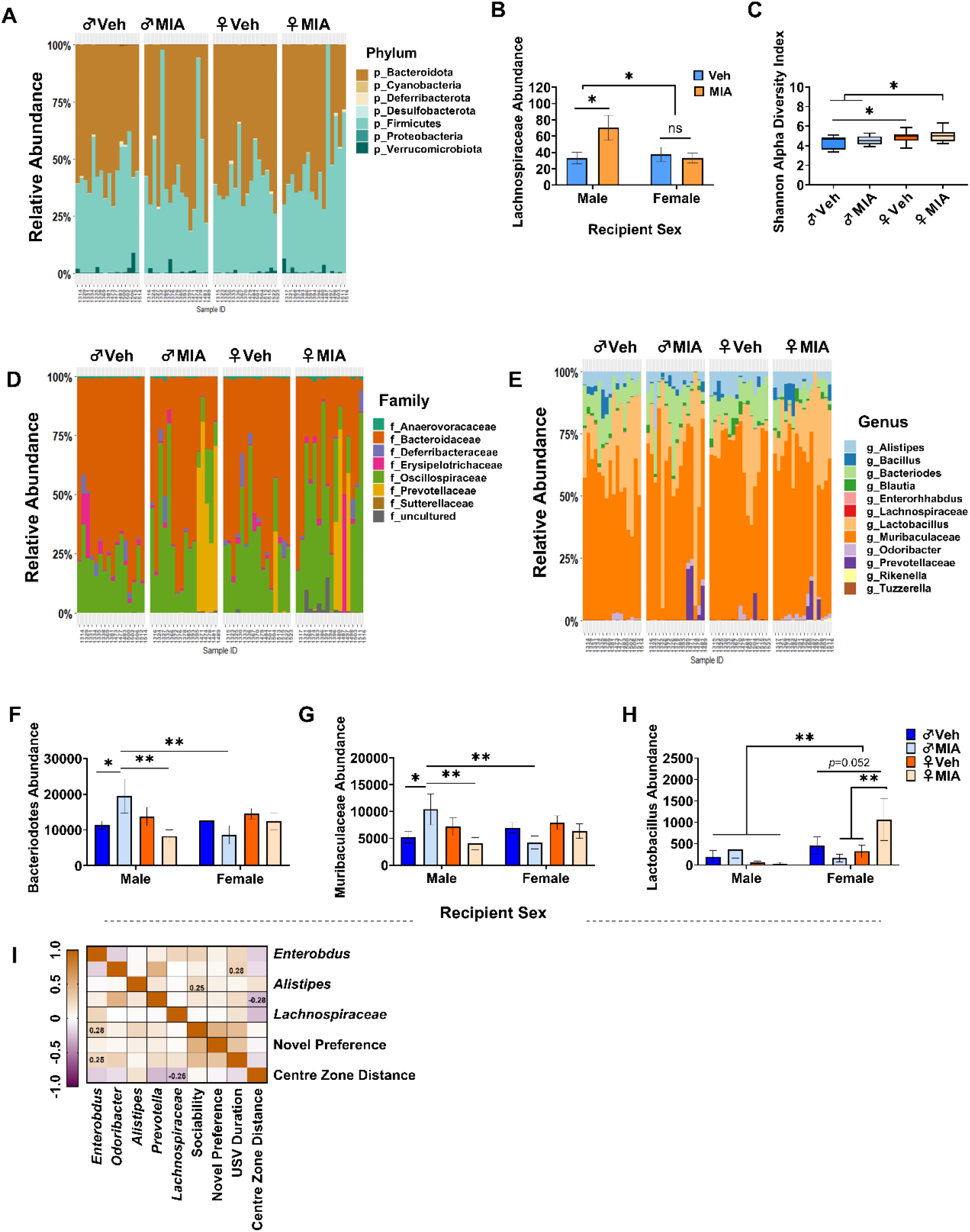
**Note:** MIA microbiota from male and female donors distinctively alters the gut composition and bacterial diversity in recipient mice. (A) composition of the gut microbiota at the phylum level in cecal samples of recipient mice. (B) MIA microbiota increased the abundance of Lachnospiraceae in male recipient mice. (C) male same-sex microbiota transplant decreased bacterial diversity. (D-E) microbiota profile at the family and genus levels in recipient mice. (F-G) male MIA microbiota increases the relative abundance of Bacteriodetes and Muribaculaceae. (H) Lactobacillus genera are increased with female MIA microbiota transplant. (I) Spearman’s rank correlation between recipient behavior and selected gut microbes. Color scale denotes Spearman’s r from brown (positive correlation) to purple (negative correlation). Data represent the mean ± SEM (*n* = 8 per group; \**p* < 0.05, \*\**p* < 0.01; n.s., not significant).

**Table 1.** Mean ± SEM of relative abundance of bacterial taxa in recipient mice treated with vehicle or MIA microbiota, regardless of donor sex.

| Taxa (phylum level) | Vehicle microbiota | MIA microbiota | *p*-value |
| --- | --- | --- | --- |
| *p_Deferribacterota* | 48.81 ± 12.55 | 14.63 ± 4.86 | **0.014** |
| *p_Proteobacteria* | 5.31 ± 2.53 | 18.13 ± 6.04 | 0.055 |
| *p_Actinobacteriota* | 38.4 ± 5.44 | 24.9 ± 4.32 | 0.056 |
| Taxa (family level) |  |  |  |
| *f_Prevotellaceae* | 51.88 ± 51.297 | 608.91 ± 260.004 | **0.038** |
| *f_Butyricicoccaceae* | N.D | 1.47 ± 0.444 | **0.002** |
| *f_Sutterellaceae* | N.D | 1.03 ± 0.452 | **0.028** |
| *f_Ruminococcaceae* | 29.9 ± 4.76 | 18.3 ± 2.76 | **0.040** |
| *f_Deferribacteraceae* | 48.81 ± 12.548 | 14.63 ± 4.862 | **0.015** |
| *f_Erysipelotrichaceae* | 59.38 ± 19.709 | 16.88 ± 2.928 | **0.037** |
| *f_Anaerovoracaceae* | 15.66 ± 1.582 | 10.44 ± 1.639 | **0.025** |
| Taxa (genus level) |  |  |  |
| *g_Prevotellaceae* | 51.875 ± 51.297 | 586.469 ± 249.288 | **0.039** |
| *g_Odoribacter* | 22.7 ± 11.9 | 70.3 ± 15.7 | **0.020** |
| *g_Rikenella* | N.D | 6.531 ± 2.331 | **0.003** |
| *g_Blautia* | 39.0 ± 8.83 | 17.0 ± 23.84 | **0.028** |
| *g_Tuzzerella* | 9.719 ± 1.958 | 4.938 ± 0.849 | **0.025** |
| *g_Parasutterella* | N.D | 1.031 ± 0.452 | **0.028** |
| *g_Lachnospiraceae* | 2.625 ± 0.619 | 0.656 ± 0.326 | **0.007** |
| *g_Eubacterium* | 92.469 ± 16.71 | 50.5 ± 9.594 | **0.032** |
| *g_Mucispirillum* | 48.813 ± 12.548 | 14.625 ± 4.862 | **0.015** |

N.D = not detected

### Male MIA microbiota transplant reduces microbiota diversity, whereas female MIA microbiota led to a differential bacterial profile that may be protective

Next, we assessed microbiota donor sex (i.e., ♂ or ♀) by treatment (i.e., Veh or MIA) interaction effects on overall microbiota diversity and abundance. The Shannon alpha diversity rarefaction index revealed that compared to ♂Veh, ♀Veh microbiota transplant led to a higher bacterial diversity in recipient mice [*H* (3) = 8.13, *p* = 0.043; *p* < 0.05 after pairwise comparison; Fig. 2C]. Such sex differences have been reported in rodents’ microbiota diversity (19), which may have a sex-differential effect. In line with this, the bacterial diversity of mice that received ♀MIA microbiota was higher than that of mice that received ♂Veh and ♂MIA microbiota (*p* < 0.05; Fig. 2C), suggesting an overall lower bacterial diversity in male donor microbiota, which is further exacerbated via MIA treatment.

Taxonomic analyses further revealed that, regardless of recipient sex, mice that received ♀MIA microbiota led to a distinct microbial community (Fig. 2D – 2E). At the family and genus levels, the abundances of *Bacteroidaceae, Enterorhabdus*, and *Bacteroides* were significantly decreased with ♀MIA microbiota transplant, whereas the abundances of *Odoribacter, Alistipes, Rikenella* which have been reported to have anti-inflammatory protective effects (29,30) were significantly increased in mice that received ♀MIA microbiota (all p values < 0.05; Table 2). The abundance of *Prevotellaceae* was, however, high in recipients of ♂MIA microbiota (*p* < 0.05; Table 2). Moreover, male mice that received ♂MIA microbiota showed an increased abundance of *Bacteroidetes* [*F* (3, 56) = 3.51, *p* = 0.021] and *Muribaculaceae* [*F* (3, 56) = 3.35, *p* = 0.025] which were rather decreased in female recipients of ♂MIA (*p* < 0.05; Fig. 2F-2G). In contrast, *Lactobacillus* genera, known for health-positive effects (31), were more abundant in female mice that received ♀MIA microbiota transplant compared to females that received ♂Veh, ♂MIA, or ♀Veh and all male recipients (*p* < 0.05; Fig. 2H), suggesting that female microbiota in female recipients may provide additional benefits. Overall, the female-specific MIA microbial profile suggests that some taxa may be protective (e.g., *Rikenella* and *Lactobacillus*), which may explain why ♂MIA, compared to ♀MIA, is more detrimental to social behavior (Fig. 1B-1G).

**Table 2.** Mean ± SEM of relative abundance of bacterial taxa in recipient mice treated with male or female vehicle or MIA microbiota microbiota.

| Taxa (family level) | ♂VEH | ♂MIA | ♀VEH | ♀MIA | p-value |
| --- | --- | --- | --- | --- | --- |
| *f__Bacteroidaceae* | 1703 ± 173.1 | 1758 ± 554.5 | 2015 ± 294.7 | 725 ± 174.5 | **0.046** |
| *f__Oscillospiraceae* | 499 ± 66.2 | 589 ± 105.9 | 858 ± 112.3 | 589 ± 77.2 | **0.046** |
| Taxa (genus level) |  |  |  |  |  |
| *g__Enterorhabdus* | 17.88 ± 3.63 | 18.50 ± 4.34 | 21.25 ± 3.58 | 7.75 ± 1.42 | **0.038** |
| *g__Bacteroides* | 1703 ± 173 | 1758 ± 554 | 2015 ± 295 | 725 ± 174 | **0.046** |
| *g__Odoribacter* | 35.81 ± 21.62 | 47.81 ± 21.75 | 9.63 ± 9.63 | 92.69 ± 21.86 | **0.030** |
| *g__Alistipes* | 34.90 ± 16.50 | 21.70 ± 11.40 | 52.80 ± 30.90 | 123.6 ± 23.50 | **0.008** |
| *g__Rikenella* | N.D | 0.813 ± 0.557 | N.D | 12.250 ± 4.218 | **< 0.001** |
| *g__Acetatifactor* | 13.00 ± 3.43 | 17.00 ± 3.52 | 59.56 ± 21.81 | 22.68 ± 6.55 | **0.024** |
| *g__Blautia* | 28.06 ± 8.23 | 23.12 ± 7.19 | 50.00 ± 15.43 | 10.87 ± 2.06 | **0.040** |
| *g__Tuzzerella* | 6.43 ± 1.27 | 5.93 ± 1.33 | 13.00 ± 3.57 | 3.93 ± 1.03 | **0.019** |
| *g__Prevotellaceae* | N.D | 880 ± 463 | 104 ± 103 | 293 ± 177 | 0.075 |

N.D = not detected

We next explored whether specific bacterial taxa correlated with the observed behavioral changes in both male and female recipient mice. The Spearman’s rank correlation revealed that four bacterial taxa co-varied with social behavior, USV average call duration, and distance travelled in the center zone of the open field test (Fig. 2I). The abundance *Enterorhabdus* and *Alistipes* (found significantly lower in ♀MIA) recipient mice, positively associated with increased social behavior and average USV call duration during social interaction with a female stimulus (all *p* values < 0.05). *Lachnospiraceae* (found significantly lower in ♀MIA recipient mice) and *Prevotella* (found significantly higher in ♂MIA recipient mice) negatively correlated with decreased anxiety-like behavior (all *p* values < 0.05; Fig. 2I). The association between distinct bacterial taxa and ASD-relevant behaviors supports previous reports that specific bacteria might play a role in diverse ASD symptoms and may be candidate targets for treatment (16,32).

### Serum cytokines and microglia changes following MIA microbiota transplantation

Alterations in the gut microbiota can impact the brain via immune functioning, and many cytokines are dysregulated in MIA offspring (33); thus, we hypothesized that MIA microbiota transplant also affects peripheral and neuroimmune factors in recipient mice. As such, we analyzed serum cytokines involved in immune function in MIA and Veh microbiota recipient mice (see SI Table 2). Overall, female recipient mice, regardless of treatment, had an increased concentration of macrophage inflammatory protein (MIP)-2 than males [*F* (1, 37) = 14.98, *p* < 0.001; Fig. 3A]. MIP-2 is a chemokine that modulates inflammatory response by recruiting neutrophils to sites of injury or infection (34), consistent with the idea that females may have a greater response to inflammatory insults (35). MIP-2 levels were also elevated in mice that received microbiota from male donors, regardless of treatment, whereas female microbiota transplant increased IL-1β in recipient mice (all *p* values < 0.05; Fig. 3B-3C). Tumor necrosis factor-alpha (TNF-α), which is altered in MIA-exposed (34), was significantly lower in female mice that received female, but not male, microbiota transplant (*p* < 0.05; Fig. 3D), suggesting an interaction between microbiota donor and recipient sex. Significant cecum treatment effects, regardless of microbiota donor sex, were also observed, with elevated levels of IL-4 [*F* (1, 37) = 4.73, *p* = 0.036] and IL-7 [*F* (1, 37) = 4.64, *p* = 0.038] in recipients of MIA microbiota compared to vehicle microbiota (*p* < 0.05; Fig. 3E-3F). IL-4 was found to correlate positively with the abundance of *Rikenella, Prevotella, Odoribacter,* and *Parasutterella* genera (all *p* values < 0.05; Fig. 3G), although no association was found between behavior and cytokine expression.

**Figure 3.**
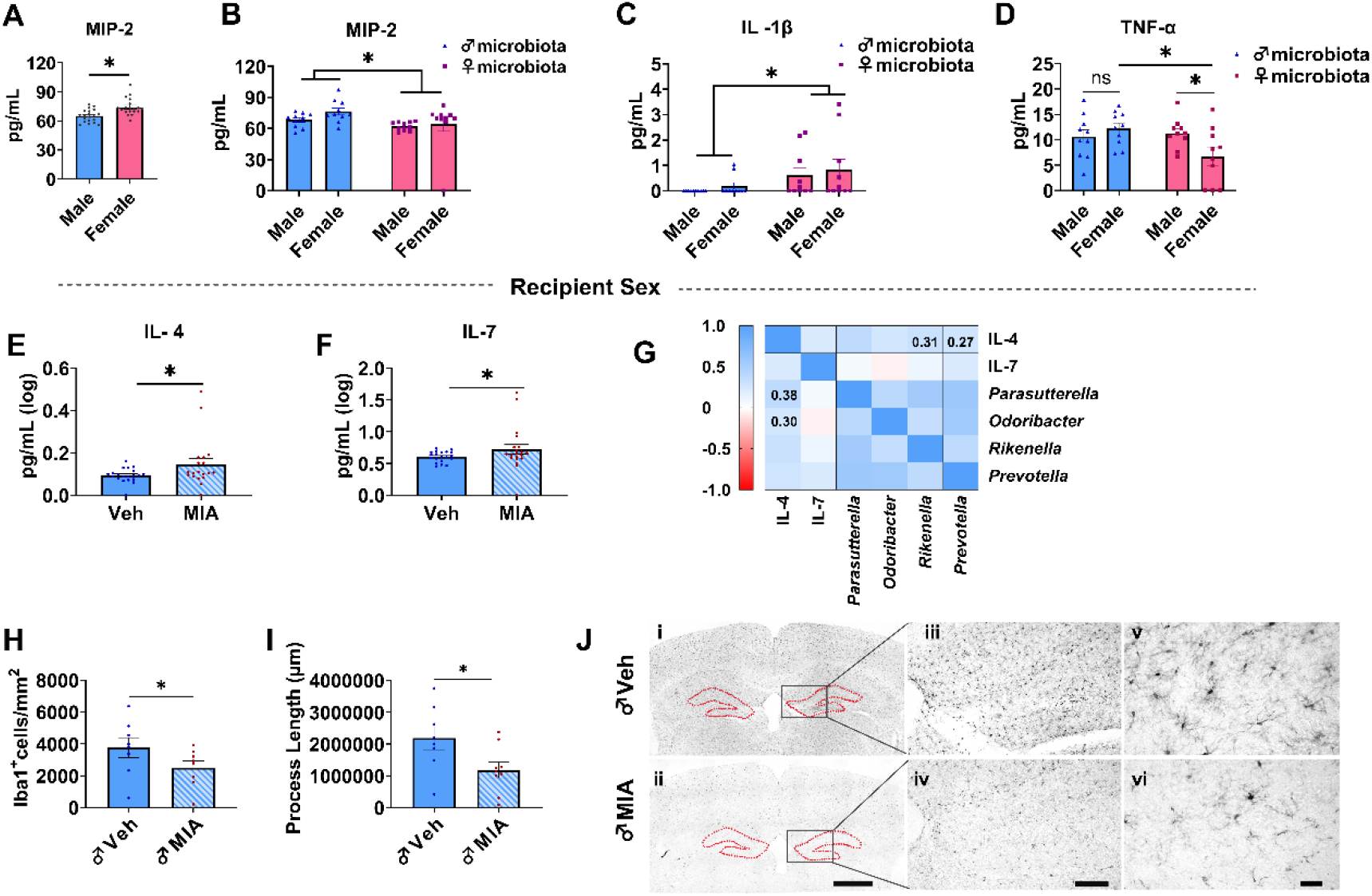
**Note:** Serum cytokines and microglia changes following MIA microbiota transplantation. (A) Recipient sex main effect. Elevated MIP-2 in females vs male recipient mice. (B & C) Donor sex main effect. Female donor microbiota decreased MIP-2 while increasing IL-1β in both sexes. (D) Recipient sex by donor sex interactions. Female donor microbiota decreased TNF-α levels in female recipient mice. (E-F) MIA microbiota increased IL-4 and IL-7 levels in recipient mice. (G) Spearman’s rank correlation between select cytokines and gut microbes. Color scale denotes Spearman’s r from purple (positive correlation) to red (negative correlation). (H-I) Male MIA microbiota decreased Iba1^+^ cells and process length in male recipient mice. Data represent the mean ± SEM (*n* = 5 per group; \**p* < 0.05, \*\**p* < 0.01; n.s., not significant). Scale bar: 300µm (i-ii); 150µm (iii-iv); 50µm (v-vi).

We next explored potential alterations in microglia activation in the hippocampal DG, a brain structure known to have a high density of microglia and strongly associated with atypical behavior (36). Using stereology, we quantified anti-ionized calcium-binding adaptor protein-1 (Iba1)+ cell numbers and morphological changes, including soma size and process length among male recipient mice with increased social behavioral deficits (Fig. 1B-1C). We found that, while microbiota from either Veh or MIA-treated female donors revealed no change in the analyzed parameters (all p values > 0.100; SI Table 3), ♂MIA microbiota transplant significantly decreased Iba1^+^ cell numbers [*F* (1, 13) = 7.05, *p* = 0.020] and process lengths [*F* (1, 13) = 8.74, *p* = 0.011] in male recipient mice (all *p* values < 0.05; Fig. 3H-3J). The reduction in cell number and microglial processes may in part, reflect an amoeboid phenotype, associated with increased phagocytosis and neuronal death (37). Together, these results suggest gut microbial changes associated with MIA may mediate ASD-like behavioral deficits via immune dysregulation.

## Discussion

Sex differences are evident in the prevalence and presentation of ASD (6,7); yet the role of biological sex is often overlooked in preclinical studies assessing sex-differential risk factors, including gut microbiota, for ASD (18–20,38). Here, we show that the gut microbiota contributes to sex differential risk and expression of ASD-like traits in the MIA mouse model of ASD. Specifically, male donor MIA microbiota induced a core ASD-like phenotype in recipients with more pronounced deficits in male recipient mice (11,12). We also find sex-specific alterations in the gut microbiota, circulating cytokines and chemokines as well as in microglial phenotype in the DG of the hippocampus. While colonization with male and female MIA microbiota induce many similar changes in gut composition and immune response, there are several indications of increased pathogenesis in recipients of male microbes and/or protective factors in recipients of female gut contents, indicating that it is the guts’ responsiveness to environmental insults, such as MIA, that contribute to the sex bias in ASD development.

In the present study, we found that MIA microbiota from both male and female donors altered the microbiota composition in a sex-specific manner. In line with previous reports (18), our male but not female donor MIA microbiota decreased bacterial diversity in recipient mice. While diversity was overall decreased, the abundance of particular gut microbes with known pathological effects were increased, including *Bacteroidetes, Blautia* and *Prevotellaceae*. ASD patients have previously been found to have high levels of similar gut bacteria (39), and these microbes have been linked to altered gut metabolites and neurotransmitter imbalances in ASD. For example, propionic acid short-chain fatty acid (SCFA) is produced by *Bacteroidetes* and has the affinity to cross the blood-brain barrier (40). *Blautia*, on the other hand, has a role in the synthesis of tryptophan, which acts as a precursor of serotonin (41). Serotonin and its modulators affect the central and enteric nervous systems, and evidence suggests aberrant serotonergic transmission in ASD (42). Specifically, the brains of patients with ASD have low levels of serotonin (43), but blood serotonin levels in a subpopulation of these children are elevated (44), which is highly correlated with gastrointestinal symptoms (45). Thus, the higher levels of *Blautia* could lead to more circulating serotonin to modulate ASD-like behavior in male recipient mice (17,46). Further, *Prevotellaceae* is linked to pathogenesis in a variety of disorders, including ASD (47) as increased *Prevotellaceae* levels lead to inflammation, mucosal activation, and systemic T-cell activation (48). Although there are mixed reports regarding the direction of alteration in *Prevotellaceae* abundance in ASD (32,49–51), it has been suggested that alteration in this microbe may be a diagnostic criterion for gut dysbiosis in ASD (32). Overall, this sex-specific overgrowth of pathogenic microbes may underlie the greater prevalence and severity of ASD-like phenotypes in recipients of male than female MIA gut microbiota, and the similarities with male MIA gut microbial profile with the human condition highlight the need to incorporate sex/gender considerations across preclinical and clinical research of gut microbiome mediation of ASD.

It is noteworthy that MIA microbiota from female donors also altered the recipient’s gut bacterial profile but with elevated levels of *Lactobacillus* and *Rikenella* genera, which may have protective effects in recipient mice. *Lactobacillus* bacteria of the family *Ruminococcaceae* improve intestinal barrier integrity, and *Lactobacillus*-derived probiotics have been found to reduce the adverse effects of stress in rodents and restore the gut alterations associated with high-fat diets in mothers (52–54). Further, *Rikenella* has previously been linked to resiliency following early-life stress (29) and is considered an essential probiotic for reducing inflammation in the intestines (55). Hence, these additional benefits of female-specific microbes likely protected recipient mice from MIA-associated social behavior deficits (e.g., our female recipient mice of female MIA microbiota exhibited a social preference than female recipients of male MIA microbiota) and may, in part, contribute to sex differences in the presentation of ASD-like behavioral abnormalities in the MIA model. Future studies may evaluate the efficacy of these sex-specific microbes in ameliorating MIA-associated behavioral deficits. Overall, our results support the existence of altered gut-brain physiological pathways in MIA models and suggest that the same immune insult may induce sex-specific gut dysbiosis and behavioral abnormalities depending on the animal’s sex.

Although our female recipients of both male and female donor vehicle microbiota maintained overall social behavior, they lacked a preference for social novelty compared to male recipients. This absence of preference for exploring novel over familiar stimuli may indicate impaired social cognition and interaction in females but could also simply reflect the sex differences in novelty-seeking behavior where female mice may not exhibit novelty preference to the same extent as males (56,57). Nonetheless, female recipient mice, regardless of donor sex or MIA status, demonstrated an overall increase in baseline activity in total distance travelled in the open field arena, which could be attributed to increased hyperactivity or heightened stress responses (58). In relation to ASD in humans, females might better camouflage or mask their social difficulties but still experience significant social anxiety and stress in novel social situations (59). Understanding sex differences at baseline, thus, has significant implications for neurodevelopmental disorders like ASD, which is crucial for developing tailored interventions and supports.

It is well-established that gut bacteria modulate the innate and adaptive immune systems of the host and that gut dysbiosis causes immune dysregulation (21,33,60). In the MIA model of ASD, pro-inflammatory cytokines such as IL-6, IL-1β and TNF-α are elevated (33). We only found elevated IL-4 and IL-7 levels with MIA microbiota transplant, although microbiota from female donors also increased pro-inflammatory IL-1β levels. Interestingly, an unexpected role for commensal bacteria in promoting IL-1β secretion to support intestinal barrier repair has recently been reported (61), and thus, the increase in this cytokine among females may be protective. Microbiota from female donors, regardless of MIA/Veh treatment, also decreased MIP-2 and TNF-α levels in female recipient mice compared to recipients of male microbiota, suggesting a greater pro-inflammatory immune response with male than female microbiota transfer. This is consistent with the gut microbiota profile, such that there may be potential protective immune reactivity from female compared to male gut microbial transfer.

Our data also indicate that these gut-immune reactivity differences also led to sex-specific effects in the morphology of the neural immune cells, microglia. Specifically, male-MIA microbiota altered microglial morphology in our male recipient mice, consistent with the neuroinflammation reported in ASD (62). Thus, the current study suggests that although the gut microbiota is disrupted in both recipients of MIA male and female gut microbiota, females exhibit protective factors within their gut and immune responses that may mitigate the impact of MIA on brain function.

Overall, our results suggest gut microbial changes associated with MIA mediate the sex bias in ASD-like behavioral deficits via gut-immune dysregulation. We have demonstrated that colonization with MIA microbiota is sufficient to promote an ASD-like phenotype in WT mice, and male MIA gut transfer to male recipients is most detrimental to social behavior. Our profiling of the gut microbiota and immune responses point to heightened pathogenicity among recipients of male MIA microbiota, coupled with protective elements in recipients of female gut contents. This suggests that the gut’s reactivity to environmental stressors, such as in the MIA model, plays a pivotal role in shaping the sex disparity observed in ASD development. Based on these findings, we propose that microbiota therapeutics may yield different outcomes based on the sex of the donor and recipient; thus, it is imperative that microbiota-based interventions take sex (and likely gender in the case of human trials) into account.

## Materials and Methods

### Animals

Male and female experimental mice were generated by mating C57BL/6 mice (Charles River Laboratories, QC, Canada). All pups remained with their mother (n = 6-7 litters/group) until weaning at post-natal day (PND) 22 into same-sex cages (2-4 mice per cage) and then singly housed from PND 28. Mice were housed at a 12-hour light-dark cycle (7 am – 7 pm) and received *ad libitum* access to food (Teklad Global Rodent Diets, Envigo) and water. All animal use and experiments adhered to the Canadian Council on Animal Care (CCAC) guidelines, approved by the Memorial University of Newfoundland and Labrador’s Institutional Animal Care Committee.

### MIA and Donor Microbiota Collection

Juvenile mice (PND 35-42) born to either poly I:C or vehicle-treated mothers served as donor mice for microbiota transplantation. Briefly, pregnant C57BL/6 mice were injected intraperitoneally on gestational day (GD) 12.5 with saline (0.9% NaCl) or 5mg/kg poly I:C (sodium salt; Sigma-Aldrich, #P1530-100MG) (13). GD 12.5 was chosen as it corresponds to the first trimester of pregnancy in humans when maternal viral infections have been linked to an increased incidence of ASD (63) with increased male susceptibility (14). Here, the cecum was excised from 6-8 MIA or vehicle-treated offspring (2*2 sexes from every litter), and contents were extracted into sterile 2-ml microcentrifuge tubes. Sex by treatment cecal contents were homogenized with 1 × phosphate-buffered saline (PBS) in a 1:1 ratio. The resulting slurry was stored at -80◦C in aliquots until the day of transplantation (i.e., for a maximum of two months between cecal microbiota collection and transplantation).

### Cecal Microbiota Transplantation

On the day of inoculation, thawed aliquots of cecal slurry were diluted further with 1× PBS in a 1:5 ratio, for a final dilution of 1:10, and vortexed gently for the supernatant isolation. Using a same-sex by cross-sex approach, experimental male and female mice (4-weeks old) were colonized with either ♂Vehicle [male (n = 12); female (n =12)], ♀ Vehicle [male (n = 12); female (n =11)], ♂ MIA [male (n = 12); female (n =12)] or ♀ MIA [male (n = 13); female (n =12)]. Each mouse in each group was colonized with 200 μL of cecal content via oral gavage every other day for 12 days. Following a 1-day rest period after oral gavage, all mice (PND 40 - 42) underwent a battery of behavioral assays to assess ASD-associated phenotype.

### Behavioral batteries

Multiple behavioral assays were selected to assess ASD core and associated relevant behaviors (i.e., social behavior, communication, restrictive and repetitive behaviors, and anxiety-like behaviors). Tests were organized in the following sequence: USVs (day 1), three-chamber social test (day 2), marble bury (day 3), and open-field tests (day 4). Mice were tested between 9am and 5pm.

### Interaction-induced juvenile USV recording and analysis

Mice emit USVs during social interaction to communicate with same-sex and opposite-sex conspecifics (22). We measured juvenile USV production in response to an unfamiliar age-matched male and female conspecific as described in (64) to assess for communication deficits. Experimental animals were placed in a recording cage (42 × 25 × 20 cm) for a 10-min habituation period, following which stimulus mice were introduced. Using an ultrasound microphone (Echo Meter Touch 2 Pro for IOS; Wildlife Acoustics attached to a 5th generation iPad running Echo Meter Touch application, version 2.8.3) suspended above the cage, we recorded USVs generated by resident mice during 3 min interactions with same-sex and opposite-sex conspecifics. The USV files were scored between 40-125 kilohertz (kHz) using spectrogram in Raven Lite software (version 2.0.4) with a scale increase of 50 ms. Calls were then analyzed for total number, total call duration (s), average call duration (s), average low frequency, and average high frequencies (kHz).

### Three-chamber social test

We performed the three-chamber test to assess sociability and preference for social novelty, as described in (65). Mice were habituated for 10 min to a 50 x 75 cm apparatus with three chambers that contained empty interaction holding cups. In the sociability test, a novel mouse of the same age and sex was placed in one of the holding cups. The testing mouse was allowed 10 min to explore the social stimulus (age and sex-matched unfamiliar mouse) vs. the empty holding cup (non-social). The location of the stimuli in either side chamber was counterbalanced between tests. An unfamiliar stimulus mouse was introduced to a previously empty holding cup in the final 10-minute session to assess for social novelty preference. Mice were then allowed to explore the now familiar mouse (stimulus mouse from previous sociability session) against the novel, unknown stimulus mouse. The time spent exploring each chamber and sniffing each interaction holding cup was recorded and analyzed by an investigator blinded to the experimental groups using automated ANY-Maze software (Stoelting Europe, Dublin, Ireland).

### Marble burying

Marble burying elicits repetitive behavior in rodents analogous to those observed in ASD (65). In a standard-sized mouse cage containing 4 cm of bedding, 16 marbles were aligned equidistantly in a 4 x 4 grid. Mice were then placed in the testing cage, and repetitive digging behaviour was quantified by the number of marbles (i.e., at least 2/3 covered in bedding) at 5-minute intervals in a 30-minute trial.

### Open field test

The open-field test assesses locomotor activity and anxiety-like behavior while exploring a novel environment (25). Prior to testing, mice were habituated to the testing room for half an hour. Mice were then placed in an open arena, and their spontaneous locomotor activity was video-recorded for 10 minutes. The total distance travelled (m) and entries into the central and outer zones were analyzed using ANY-Maze.

### Dissections

Upon completion of behavior testing, all experimental mice (PND 44-46) were sacrificed by rapid decapitation under injectable anesthesia (Avertin 40mg/kg) for blood and tissue collection. Blood was stored a 4 ◦C for 24 hours, then centrifuged (3000 rcf × 8 °C for 20 min) for serum isolation and stored at −80°C until further processing. Brains were extracted and post-fixed for 4 hours in 4% paraformaldehyde and then transferred to 20% sucrose, where they were stored at 4 ◦C until sectioning with the microtome. Serial 30μm free-floating sections were collected and stored at −20 °C in cryoprotectant until use. Cecal contents were also collected in Eppendorf tubes and stored at −80 ◦C until prepared for transfer for bacterial sequencing.

### 16S sRNA sequencing analysis

For gene sequencing and bioinformatics, frozen cecal samples were shipped on dry ice to Integrated Microbiome Resource (Dalhousie University, Halifax). Bacterial gene amplicons were generated using the new Microbiome Helper pipeline based on the open-source pipeline Quantitative Insights into Microbial Ecology version 2 (QIIME 2.0) (66) and subsequently normalized for analysis. Following quality filtering, 1,379,130 features were retained in 64 samples (8 samples x 8 groups), averaging 21,548.90 and ranging from 3,139 - 65,335. Alpha diversity estimates (Observed Species and Shannon’s diversity) were calculated and compared between groups using the Kruskal-Wallis nonparametric test (*p* < 0.05). Additionally, changes in microbial composition between different experimental groups were assessed at each taxonomic level.

### Serum cytokine profiling

Frozen serum samples (*n*=5 per group) were shipped on dry ice to Eve Technologies (Calgary, Alberta) for multiplexed mouse high-sensitivity 18-Plex Discovery Assay® (MilliporeSigma, Burlington, Massachusetts, USA), following the manufacturer’s protocol. This 18-plex included GM-CSF, IFNγ, IL-1α, IL-1β, IL-2, IL-4, IL-5, IL-6, IL-7, IL-10, IL-12(p70), IL-13, IL-17A, KC/CXCL1, LIX, MCP-1, MIP-2, and TNFα. The assay sensitivities for these markers range from 0.06 to 9.06 pg/mL within the 18-plex panel. Individual analyte sensitivity values are available in the MilliporeSigma MILLIPLEX® MAP protocol.

### Iba1 Immunohistochemistry

A random selection of serial free-floating brain sections of experimental male recipient mice (*n*=5 per group) was used for Iba1 immunohistochemistry analysis. First, sections were washed thrice in Tris buffer before being incubated in 1% hydrogen peroxide (H2O2) for 30 minutes. Sections were then washed and blocked for 1 hour at room temperature in 10% normal goat serum (0.1% Triton X in Tris buffer; Vector Labs, Cat# S2000). Both primary and secondary antibodies were diluted in the same Tris buffer. Following two 10-minute washes, sections were incubated in primary antibody (1:10,000; Iba1 anti-rabbit, Cedarlane Labs, Cat# 019-19741) for 60 hours on a rotator with ice on, rinsed with Tris buffer, and then incubated in secondary antibody (1:1000; goat anti-rabbit IgG, Vector Labs Cat# BA-1000) for an hour at room temperature. Sections were washed and incubated in ABC Elite Kit (1:1000, Vector Labs Cat# PK6100) for 1.5 h, washed and then incubated in Metal Enhanced DAB Substate, according to manufacturers protocol (Thermo Scientific Cat# 34065) for approximately 8 minutes and mounted on gelatin-subbed slides for microscopic analyses.

### Unbiased stereological quantification of Iba1 microglia

An unbiased stereological assessment was conducted using a Leica brightfield microscope with a three-axis motorized stage, camera, and computer with Stereologer software by an investigator blinded to the experimental groups. The bilateral hippocampal DG region of interest (ROI) was outlined at 5x objective according to the atlas of (67), and Iba1 labelled cells were counted using a 60x oil objective lens. The cell body (soma) was used as the counting target for Iba1+ cells, and their numbers were estimated using the optical fractionator method. For analysis, we set an optical dissector height of 10μm with 2μm guard zone on top and bottom and used a 50 x 50 μm counting frame throughout each cell section. Total number of sections with DG was approximated by the program, taking into account number of series and number of sections with DG within the series underanalysis. We analyzed 3 sections of the DG for each animal. We then used the Cavalieri estimator to approximate whole DG and cell body volumes (μm3) for each microglia cell counted. Microglial process lengths were determined with a Spaceballs probe diameter of 6 μm.

### Statistical analysis and data availability

All statistical analyses were performed using Jamovi software (version 2.2.5; Open-source Project., Sydney, Australia) at the 0.05 alpha significance level. Most datasets (behavior, microbiota composition, serum cytokine concentration, and Iba1+ microglia morphology) were analyzed with two-way analysis of variance (ANOVA) with microbiota donor sex (♂ vs ♀) or treatment (Vehicle vs MIA) and recipient sex (male vs female) as between factors followed by Fisher’s LSD post-hoc test on significant interactions. The repeated measures ANOVA was also performed with chamber/zones as within factors on the three-chamber data. Spearman’s correlations were applied to bacterial taxa, inflammatory markers, and behavioral parameters. Graphs were generated in GraphPad Prism (version 10.2.3), and data are presented as mean ± SEM with individual datapoints depicted over bars.

## Acknowledgments

We thank the animal care services at the Memorial University of Newfoundland for their assistance with animal husbandry. Research was supported by a Discovery Grant from the Natural Sciences and Engineering Council of Canada (NSERC) and Canadian Institutes of Health Research (CIHR) Project Grant to AS-G.

## Author Contributions

S.S. and A.S.G. designed research; S.S., F.F.B., M.H., S.G.W. and A.M.R. performed research; M.A.M. performed stereology analyses on Iba1+ cells; A.S.G. supervised; S.S. and A.S.G. analyzed data; S.S. wrote the paper and review/editing by S.S., A.S.G., S.G.W., F.F.B., M.H., A.M.R., and M.A.M.

## Competing Interest Statement

The authors disclose no competing interests.

## Supplementary tables

For supplementary tables 1-3, means and SEM (standard error of the mean) are presented for the effects of same-sex and cross-sex MIA or vehicle microbiota transplantation on juvenile ultrasonic vocations (USVs), microglia alteration, and cytokine profile in recipient mice.

**Table S1.** Means ± SEM for juvenile USV parameters in recipient male and female mice interacting with a female conspecific.

| USV parameter | Male Recipients<br>(*n* = 49) | Female Recipients<br>(*n* = 47) |
| --- | --- | --- |
| Number of Calls | 476.20 ± 89.02 | 27.53 ± 19.56 |
| Average Call Duration (s) | 0.02 ± 0.00 | 0.01 ± 0.00 |
| Total Call Duration (s) | 11.89 ± 2.32 | 0.69 ± 0.52 |
| Average Total Call Frequency (kHz) | 59917.80 ± 3917.11 | 31581.09 ± 5048.34 |

**Table S2:** Means ± SEM for serum cytokine concentration in male and female recipient mice by cecum sex/treatment.

|  | ♂Veh |  | ♂MIA |  | ♀Veh |  | ♀MIA |  |
| --- | --- | --- | --- | --- | --- | --- | --- | --- |
| Cytokine (pg/mL) | Male (n = 5) | Female (n = 5) | Male (n = 5) | Female (n = 5) | Male (n = 5) | Female (n = 5) | Male (n = 5) | Female (n = 4) |
| IL-1β | N.D | N.D | N.D | 0.38 ± 0.24 | 0.46 ± 0.46 | 0.92 ± 0.76 | 0.78 ± 0.40 | 1.37 ± 1.01 |
| IL-4 | 0.09 ± 0.03 | 0.09 ± 0.01 | 0.19 ± 0.06 | 0.10 ± 0.01 | 0.10 ± 0.01 | 0.09 ± 0.01 | 0.19 ± 0.08 | 0.32 ± 0.19 |
| IL-6 | 3.54 ± 0.33 | 3.59 ± 1.01 | 4.26 ± 0.94 | 3.47 ± 0.29 | 3.33 ± 0.13 | 3.24 ± 0.50 | 3.56 ± 0.39 | 6.83 ± 3.29 |
| IL-7 | 0.59 ± 0.03 | 0.59 ± 0.04 | 0.86 ± 0.18 | 0.68 ± 0.08 | 0.64 ± 0.04 | 0.61 ± 0.04 | 0.85 ± 0.20 | 0.64 ± 0.02 |
| TNF-α | 11.58 ± 1.84 | 11.43 ± 1.68 | 9.68 ± 2.14 | 13.04 ± 1.37 | 10.24 ± 0.96 | 6.37 ± 2.03 | 12.30 ± 1.57 | 8.75 ± 3.32 |
| MIP-2 | 68.14 ± 3.46 | 78.77 ± 4.76 | 68.38 ± 3.21 | 74.16 ± 5.18 | 60.55 ± 1.61 | 72.96 ± 2.69 | 63.76 ± 2.04 | 70.39 ± 0.78 |

**Table S3:** Means ± SEM for microglia density parameters in male recipient mice by cecum sex/treatment.

|  | ♂Veh<br>(*n* = 7) | ♂MIA<br>(*n* = 8) | ♀Veh<br>(*n* = 5) | ♀MIA<br>(*n* = 5) |
| --- | --- | --- | --- | --- |
| Cell Count | 4221.00 ± 490.14 | 2505.00 ± 426.63 | 4327.80 ± 453.57 | 3949.90 ± 577.79 |
| Soma Size ($\mu^3$) | 120.00 ± 10.41 | 104.00 ± 9.92 | 108.50 ± 13.09 | 96.10 ± 8.72 |
| Space Ball (Process length) ($\mu$) | 2440.00 ± 299.60 | 1290.00 ± 251.02 | 2350.00 ± 376.52 | 2280.00 ± 404.05 |
| DG ($\mu^3$) | 157000.00 ± 10800.00 | 143000.00 ± 11000.00 | 139000.00 ± 10200.00 | 131000.00 ± 17900.00 |

## Notes

### Competing Interest Statement

The authors have declared no competing interest.

## References

1. Lord, C., Elsabbagh, M., Baird, G., & Veenstra-Vanderweele, J. (2018). Autism spectrum disorder. Lancet (London, England), 392(10146), 508–520. 10.1016/S0140-6736(18)31129-2

2. Zeidan, J., Fombonne, E., Scorah, J., Ibrahim, A., Durkin, M. S., Saxena, S., Yusuf, A., Shih, A., & Elsabbagh, M. (2022). Global prevalence of autism: A systematic review update. Autism research: Official Journal of the International Society for Autism Research, 15(5), 778–790. 10.1002/aur.2696

3. Rogge, N., & Janssen, J. (2019). The economic costs of autism spectrum disorder: A literature review. Journal of Autism and Developmental Disorders, 49(7), 2873–2900. 10.1007/s10803-019-04014-z

4. Drapeau, E., Riad, M., Kajiwara, Y., & Buxbaum, J. D. (2018). Behavioral phenotyping of an improved mouse model of phelan-mcdermid syndrome with a complete deletion of the Shank3 gene. eNeuro, 5(3), ENEURO.0046-18.2018. 10.1523/ENEURO.0046-18.2018

5. Loomes, R., Hull, L., & Mandy, W. P. L. (2017). What is the male-to-female ratio in autism spectrum disorder? A systematic review and meta-analysis. Journal of the American Academy of Child and Adolescent Psychiatry, 56(6), 466–474. 10.1016/j.jaac.2017.03.013

6. Werling, D. M., & Geschwind, D. H. (2013). Sex differences in autism spectrum disorders. Current Opinion in Neurology, 26(2), 146–153. 10.1097/WCO.0b013e32835ee548

7. Werling D. M. (2016). The role of sex-differential biology in risk for autism spectrum disorder. Biology of Sex Differences, 7, 58. 10.1186/s13293-016-0112-8

8. Patel, S., Dale, R. C., Rose, D., Heath, B., Nordahl, C. W., Rogers, S., Guastella, A. J., & Ashwood, P. (2020). Maternal immune conditions are increased in males with autism spectrum disorders and are associated with behavioural and emotional but not cognitive comorbidity. Translational Psychiatry, 10(1), 286. 10.1038/s41398-020-00976-2

9. Choi, G. B., Yim, Y. S., Wong, H., Kim, S., Kim, H., Kim, S. V., Hoeffer, C. A., Littman, D. R., & Huh, J. R. (2016). The maternal interleukin-17a pathway in mice promotes autism-like phenotypes in offspring. Science (New York, N.Y.), 351(6276), 933–939. 10.1126/science.aad0314

10. Smith, S. E., Li, J., Garbett, K., Mirnics, K., & Patterson, P. H. (2007). Maternal immune activation alters fetal brain development through interleukin-6. The Journal of Neuroscience : The Official Journal of the Society for Neuroscience, 27(40), 10695– 10702. 10.1523/JNEUROSCI.2178-07.2007

11. Haida, O., Al Sagheer, T., Balbous, A., Francheteau, M., Matas, E., Soria, F., Fernagut, P. O., & Jaber, M. (2019). Sex-dependent behavioral deficits and neuropathology in a maternal immune activation model of autism. Translational Psychiatry, 9(1), 1–12. 10.1038/s41398-019-0457-y

12. Hsiao, E. Y., McBride, S. W., Hsien, S., Sharon, G., Hyde, E. R., McCue, T., Codelli, J. A., Chow, J., Reisman, S. E., Petrosino, J. F., Patterson, P. H., & Mazmanian, S. K. (2013). Microbiota modulate behavioral and physiological abnormalities associated with neurodevelopmental disorders. Cell, 155(7), 1451–1463.

13. Juckel, G., Manitz, M. P., Freund, N., & Gatermann, S. (2021). Impact of Poly I:C induced maternal immune activation on offspring’s gut microbiome diversity - Implications for schizophrenia. Progress in Neuro-psychopharmacology & Biological Psychiatry, 110, 110306. 10.1016/j.pnpbp.2021.110306

14. Carabotti, M., Scirocco, A., Maselli, M. A., & Severi, C. (2015). The gut-brain axis: interactions between enteric microbiota, central and enteric nervous systems. Annals of Gastroenterology, 28(2), 203–209.

15. Sandoval-Motta, S., Aldana, M., Martínez-Romero, E., & Frank, A. (2017). The human microbiome and the missing heritability problem. Frontiers in Genetics, 8, 80. 10.3389/fgene.2017.00080

16. Sharon, G., Cruz, N. J., Kang, D. W., Gandal, M. J., Wang, B., Kim, Y. M., … & Mazmanian, S. K. (2019). Human gut microbiota from autism spectrum disorder promote behavioral symptoms in mice. Cell, 177(6), 1600–1618.e17. 10.1016/j.cell.2019.05.004

17. Xiao, L., Yan, J., Yang, T., Zhu, J., Li, T., Wei, H., & Chen, J. (2021). Fecal microbiome transplantation from children with autism spectrum disorder modulates tryptophan and serotonergic synapse metabolism and induces altered behaviors in germ-free mice. Msystems, 6(2), e01343–20. 10.1128/mSystems.01343-20

18. Kim, Y. S., Unno, T., Kim, B. Y., & Park, M. S. (2020). Sex differences in gut microbiota. The World Journal of Men’s Health, 38(1), 48–60. 10.5534/wjmh.190009

19. Desbonnet, L., Clarke, G., Shanahan, F., Dinan, T. G., & Cryan, J. F. (2014). Microbiota is essential for social development in the mouse. Molecular psychiatry, 19(2), 146–148. 10.1038/mp.2013.65

20. Salia, S., Martin, Y., Burke, F. F., Myles, L. A., Jackman, L., Halievski, K., Bambico, F. R., & Swift-Gallant, A. (2023). Antibiotic-induced socio-sexual behavioral deficits are reversed via cecal microbiota transplantation but not androgen treatment. *Brain, Behavior*, & Immunity - Health, 30, 100637. 10.1016/j.bbih.2023.100637

21. Thion, M. S., Low, D., Silvin, A., Chen, J., Grisel, P., Schulte-Schrepping, J., Blecher, R., Ulas, T., Squarzoni, P., Hoeffel, G., Coulpier, F., Siopi, E., David, F. S., Scholz, C., Shihui, F., Lum, J., Amoyo, A. A., Larbi, A., Poidinger, M., Buttgereit, A., … Garel, S. (2018). Microbiome influences prenatal and adult microglia in a sex-specific manner. Cell, 172(3), 500–516.e16. 10.1016/j.cell.2017.11.042

22. Scattoni, M. L., Michetti, C., & Ricceri, L. (2018). Rodent vocalization studies in animal models of the autism spectrum disorder. In Handbook of Behavioral Neuroscience (Vol. 25, pp. 445-456). Elsevier.

23. Kisko, T. M., Euston, D. R., & Pellis, S. M. (2015). Are 50-khz calls used as play signals in the playful interactions of rats? III. The effects of devocalization on play with unfamiliar partners as juveniles and as adults. Behavioural Processes, 113, 113–121. 10.1016/j.beproc.2015.01.016

24. Thomas, A., Burant, A., Bui, N., Graham, D., Yuva-Paylor, L. A., & Paylor, R. (2009). Marble burying reflects a repetitive and perseverative behavior more than novelty-induced anxiety. Psychopharmacology, 204, 361–373.

25. Seibenhener, M. L., & Wooten, M. C. (2015). Use of the Open Field Maze to measure locomotor and anxiety-like behavior in mice. Journal of Visualized Experiments: JoVE, (96), e52434. 10.3791/52434

26. Zaboski, B. A., & Storch, E. A. (2018). Comorbid autism spectrum disorder and anxiety disorders: a brief review. Future Neurology, 13(1), 31–37.

27. Tartaglione, A. M., Villani, A., Ajmone-Cat, M. A., Minghetti, L., Ricceri, L., Pazienza, V., De Simone, R., & Calamandrei, G. (2022). Maternal immune activation induces autism-like changes in behavior, neuroinflammatory profile, and gut microbiota in mouse offspring of both sexes. Translational Psychiatry, 12(1), 384. 10.1038/s41398-022-02149-9

28. Laue, H. E., Korrick, S. A., Baker, E. R., Karagas, M. R., & Madan, J. C. (2020). Prospective associations of the infant gut microbiome and microbial function with social behaviors related to autism at age 3 years. Scientific Reports, 10(1), 15515. 10.1038/s41598-020-72386-9

29. Pusceddu, M. M., El Aidy, S., Crispie, F., O’Sullivan, O., Cotter, P., Stanton, C., Kelly, P., Cryan, J. F., & Dinan, T. G. (2015). N-3 polyunsaturated fatty acids (PUFAs) reverse the impact of early-life stress on the gut microbiota. PloS One, 10(10), e0139721. 10.1371/journal.pone.0139721

30. Thu Thuy Nguyen, V., & Endres, K. (2022). Targeting gut microbiota to alleviate neuroinflammation in Alzheimer’s disease. Advanced Drug Delivery Reviews, 188, 114418. 10.1016/j.addr.2022.114418

31. He, X., Liu, W., Tang, F., Chen, X., & Song, G. (2023). Effects of probiotics on autism spectrum disorder in children: a systematic review and meta-analysis of clinical trials. Nutrients, 15(6), 1415. 10.3390/nu15061415

32. Agarwala, S., Naik, B., & Ramachandra, N. B. (2021). Mucosa-associated specific bacterial species disrupt the intestinal epithelial barrier in the autism phenome. Brain, behavior, & immunity - health, 15, 100269. 10.1016/j.bbih.2021.100269

33. Garay, P. A., Hsiao, E. Y., Patterson, P. H., & McAllister, A. K. (2013). Maternal immune activation causes age- and region-specific changes in brain cytokines in offspring throughout development. *Brain*, Behavior, and Immunity, 31, 54–68. 10.1016/j.bbi.2012.07.008

34. Qin, C. C., Liu, Y. N., Hu, Y., Yang, Y., & Chen, Z. (2017). Macrophage inflammatory protein-2 as mediator of inflammation in acute liver injury. World Journal of Gastroenterology, 23(17), 3043–3052. 10.3748/wjg.v23.i17.3043

35. Klein, S. L., & Flanagan, K. L. (2016). Sex differences in immune responses. Nature Reviews. Immunology, 16(10), 626–638. 10.1038/nri.2016.90

36. Fernández de Cossío, L., Lacabanne, C., Bordeleau, M., Castino, G., Kyriakakis, P., & Tremblay, M. È. (2021). Lipopolysaccharide-induced maternal immune activation modulates microglial CX3CR1 protein expression and morphological phenotype in the hippocampus and dentate gyrus, resulting in cognitive inflexibility during late adolescence. Brain, Behavior, and Immunity, 97, 440–454.

37. Lier, J., Streit, W. J., & Bechmann, I. (2021). Beyond activation: Characterizing microglial functional phenotypes. Cells, 10(9), 2236. 10.3390/cells10092236

38. Valeri, F., & Endres, K. (2021). How biological sex of the host shapes its gut microbiota. Frontiers in Neuroendocrinology, 61, 100912. 10.1016/j.yfrne.2021.100912

39. Wu, T., Wang, H., Lu, W., Zhai, Q., Zhang, Q., Yuan, W., Gu, Z., Zhao, J., Zhang, H., & Chen, W. (2020). Potential of gut microbiome for detection of autism spectrum disorder. Microbial Pathogenesis, 149, 104568. 10.1016/j.micpath.2020.104568

40. Thomas, R. H., Meeking, M. M., Mepham, J. R., Tichenoff, L., Possmayer, F., Liu, S., & MacFabe, D. F. (2012). The enteric bacterial metabolite propionic acid alters brain and plasma phospholipid molecular species: further development of a rodent model of autism spectrum disorders. Journal of Neuroinflammation, 9, 153. 10.1186/1742-2094-9-153

41. Golubeva, A. V., Joyce, S. A., Moloney, G., Burokas, A., Sherwin, E., Arboleya, S., Flynn, I., Khochanskiy, D., Moya-Pérez, A., Peterson, V., Rea, K., Murphy, K., Makarova, O., Buravkov, S., Hyland, N. P., Stanton, C., Clarke, G., Gahan, C. G. M., Dinan, T. G., & Cryan, J. F. (2017). Microbiota-related changes in bile acid & tryptophan metabolism are associated with gastrointestinal dysfunction in a mouse model of autism. EBioMedicine, 24, 166–178. 10.1016/j.ebiom.2017.09.020

42. Israelyan, N., & Margolis, K. G. (2019). Reprint of: Serotonin as a link between the gut-brain-microbiome axis in autism spectrum disorders. Pharmacological Research, 140, 115–120. 10.1016/j.phrs.2018.12.023

43. Adamsen, D., Ramaekers, V., Ho, H. T., Britschgi, C., Rüfenacht, V., Meili, D., Bobrowski, E., Philippe, P., Nava, C., Van Maldergem, L., Bruggmann, R., Walitza, S., Wang, J., Grünblatt, E., & Thöny, B. (2014). Autism spectrum disorder associated with low serotonin in CSF and mutations in the SLC29A4 plasma membrane monoamine transporter (PMAT) gene. Molecular Autism, 5, 43. 10.1186/2040-2392-5-43

44. Gabriele, S., Sacco, R., & Persico, A. M. (2014). Blood serotonin levels in autism spectrum disorder: a systematic review and meta-analysis. European neuropsychopharmacology : The Journal of the European College of Neuropsychopharmacology, 24(6), 919–929. 10.1016/j.euroneuro.2014.02.004

45. Marler, S., Ferguson, B. J., Lee, E. B., Peters, B., Williams, K. C., McDonnell, E., Macklin, E. A., Levitt, P., Gillespie, C. H., Anderson, G. M., Margolis, K. G., Beversdorf, D. Q., & Veenstra-VanderWeele, J. (2016). Brief Report: Whole Blood Serotonin Levels and Gastrointestinal Symptoms in Autism Spectrum Disorder. Journal of Autism and Developmental Disorders, 46(3), 1124–1130. 10.1007/s10803-015-2646-8

46. Muller, C. L., Anacker, A. M. J., & Veenstra-VanderWeele, J. (2016). The serotonin system in autism spectrum disorder: From biomarker to animal models. Neuroscience, 321, 24–41. 10.1016/j.neuroscience.2015.11.010

47. Abdelsalam, N. A., Hegazy, S. M., & Aziz, R. K. (2023). The curious case of Prevotella copri. Gut Microbes, 15(2), 2249152. 10.1080/19490976.2023.2249152

48. Iljazovic, A., Roy, U., Gálvez, E. J. C., Lesker, T. R., Zhao, B., Gronow, A., Amend, L., Will, S. E., Hofmann, J. D., Pils, M. C., Schmidt-Hohagen, K., Neumann-Schaal, M., & Strowig, T. (2021). Perturbation of the gut microbiome by Prevotella spp. enhances host susceptibility to mucosal inflammation. Mucosal Immunology, 14(1), 113–124. 10.1038/s41385-020-0296-4

49. Zou, R., Xu, F., Wang, Y., Duan, M., Guo, M., Zhang, Q., Zhao, H., & Zheng, H. (2020). Changes in the Gut Microbiota of Children with Autism Spectrum Disorder. Autism Research : Official Journal of the International Society for Autism Research, 13(9), 1614–1625. 10.1002/aur.2358

50. Morton, J. T., Jin, D. M., Mills, R. H., Shao, Y., Rahman, G., McDonald, D., Zhu, Q., Balaban, M., Jiang, Y., Cantrell, K., Gonzalez, A., Carmel, J., Frankiensztajn, L. M., Martin-Brevet, S., Berding, K., Needham, B. D., Zurita, M. F., David, M., Averina, O. V., Kovtun, A. S., … Taroncher-Oldenburg, G. (2023). Multi-level analysis of the gut-brain axis shows autism spectrum disorder-associated molecular and microbial profiles. Nature Neuroscience, 26(7), 1208–1217. 10.1038/s41593-023-01361-053

51. Kang, D. W., Adams, J. B., Coleman, D. M., Pollard, E. L., Maldonado, J., McDonough-Means, S., Caporaso, J. G., & Krajmalnik-Brown, R. (2019). Long-term benefit of Microbiota Transfer Therapy on autism symptoms and gut microbiota. Scientific Reports, 9(1), 5821. 10.1038/s41598-019-42183-0

52. He, J., Wang, W., Wu, Z., Pan, D., Guo, Y., Cai, Z., & Lian, L. (2019). Effect of Lactobacillus reuteri on intestinal microbiota and immune parameters: Involvement of sex differences. Journal of Functional Foods, 53, 36–43.

53. Mindus, C., Ellis, J., van Staaveren, N., & Harlander-Matauschek, A. (2021). Lactobacillus-based probiotics reduce the adverse effects of stress in rodents: A meta-analysis. Frontiers in Behavioral Neuroscience, 15, 642757. 10.3389/fnbeh.2021.642757

54. Buffington, S. A., Di Prisco, G. V., Auchtung, T. A., Ajami, N. J., Petrosino, J. F., & Costa-Mattioli, M. (2016). Microbial reconstitution reverses maternal diet-induced social and synaptic deficits in offspring. Cell, 165(7), 1762–1775. 10.1016/j.cell.2016.06.001

55. Cox, L. M., Yamanishi, S., Sohn, J., Alekseyenko, A. V., Leung, J. M., Cho, I., Kim, S. G., Li, H., Gao, Z., Mahana, D., Zárate Rodriguez, J. G., Rogers, A. B., Robine, N., Loke, P., & Blaser, M. J. (2014). Altering the intestinal microbiota during a critical developmental window has lasting metabolic consequences. Cell, 158(4), 705–721. 10.1016/j.cell.2014.05.052

56. Granza, A. E., Amaral, I. M., Monteiro, D. G., Salti, A., Hofer, A., & El Rawas, R. (2023). Social Interaction Is Less Rewarding in Adult Female than in Male Mice. Brain Sciences, 13(10), 1445. 10.3390/brainsci13101445

57. Kopachev, N., Netser, S., & Wagner, S. (2022). Sex-dependent features of social behavior differ between distinct laboratory mouse strains and their mixed offspring. iScience, 25(2), 103735. 10.1016/j.isci.2022.103735

58. Deal, A., Cooper, N., Kirse, H. A., Uneri, A., Raab-Graham, K., Weiner, J. L., & Solberg Woods, L. C. (2021). Early life stress induces hyperactivity but not increased anxiety-like behavior or ethanol drinking in outbred heterogeneous stock rats. *Alcohol (Fayetteville*, N.Y.), 91, 41–51. 10.1016/j.alcohol.2020.11.007

59. Belcher, H. L., Morein-Zamir, S., Stagg, S. D., & Ford, R. M. (2023). Shining a Light on a Hidden Population: Social Functioning and Mental Health in Women Reporting Autistic Traits But Lacking Diagnosis. Journal of Autism and Developmental Disorders, 53(8), 3118–3132. 10.1007/s10803-022-05583-2

60. Mazmanian, S. K., Liu, C. H., Tzianabos, A. O., & Kasper, D. L. (2005). An immunomodulatory molecule of symbiotic bacteria directs maturation of the host immune system. Cell, 122(1), 107–118. 10.1016/j.cell.2005.05.007

61. Wu, W. H., Kim, M., Chang, L. C., Assie, A., Saldana-Morales, F. B., Zegarra-Ruiz, D. F., Norwood, K., Samuel, B. S., & Diehl, G. E. (2022). Interleukin-1β secretion induced by mucosa-associated gut commensal bacteria promotes intestinal barrier repair. Gut Microbes, 14(1), 2014772. 10.1080/19490976.2021.2014772

62. Vargas, D. L., Nascimbene, C., Krishnan, C., Zimmerman, A. W., & Pardo, C. A. (2005). Neuroglial activation and neuroinflammation in the brain of patients with autism. Annals of Neurology, 57(1), 67–81. 10.1002/ana.20315

63. Atladóttir, H. O., Thorsen, P., Schendel, D. E., Østergaard, L., Lemcke, S., & Parner, E. T. (2010). Association of hospitalization for infection in childhood with diagnosis of autism spectrum disorders: a Danish cohort study. Archives of Pediatrics & Adolescent Medicine, 164(5), 470–477. 10.1001/archpediatrics.2010.9

64. Scattoni, M. L., Crawley, J., & Ricceri, L. (2009). Ultrasonic vocalizations: a tool for behavioural phenotyping of mouse models of neurodevelopmental disorders. Neuroscience and Biobehavioral Reviews, 33(4), 508–515. 10.1016/j.neubiorev.2008.08.003

65. Chang, Y. C., Cole, T. B., & Costa, L. G. (2017). Behavioral Phenotyping for Autism Spectrum Disorders in Mice. Current Protocols in Toxicology, 72, 11.22.1–11.22.21. 10.1002/cptx.19

66. Bolyen, E., Rideout, J.R., Dillon, M.R., Bokulich, N.A., Abnet, C.C., Al-Ghalith, G.A., et al., 2019. Reproducible, interactive, scalable and extensible microbiome data science using QIIME 2. Nature Biotechnology. 37, 852–857. 10.1038/ s41587-019-0209-9.

67. Paxinos, G. and Franklin, K.B.J. (2001) The Mouse Brain in Stereotaxic Coordinates. 2nd Edition, Academic Press, San Diego.

